# Properties of rhythmogenic currents in spinal Shox2 interneurons across postnatal development

**DOI:** 10.1101/2024.09.26.612677

**Authors:** Shayna Singh, Lihua Yao, Natalia A. Shevtsova, Ilya A. Rybak, Kimberly J. Dougherty

## Abstract

Locomotor behaviors are performed by organisms throughout life, despite developmental changes in cellular properties, neural connectivity, and biomechanics. The basic rhythmic activity in the central nervous system that underlies locomotion is thought to be generated via a complex balance between network and intrinsic cellular properties. Within mature mammalian spinal locomotor circuitry, we have yet to determine which properties of spinal interneurons (INs) are critical to rhythmogenesis and how they change during development. Here, we combined whole cell patch clamp recordings, immunohistochemistry, and RNAscope targeting lumbar Shox2 INs in mice, which are known to be involved in locomotor rhythm generation. We focused on the properties of putatively rhythmogenic ionic currents and the expression of corresponding ion channels across postnatal time points in mice. We show that subsets of Shox2 INs display voltage-sensitive conductances, in addition to respective ion channels, which may contribute to or shape rhythmic bursting. Persistent inward currents, M-type potassium currents, slow afterhyperpolarization, and T-type calcium currents are enhanced with age. In contrast, the hyperpolarization-activated and A-type potassium currents were either found with low prevalence in subsets of neonatal, juvenile, and adult Shox2 INs or did not developmentally change. We show that Shox2 INs become more electrophysiologically diverse by juvenile and adult ages, when locomotor behavior is weight-bearing. These results suggest a developmental shift in the magnitude of rhythmogenic ionic currents and the expression of corresponding ion channels that may be important for mature, weight-bearing locomotor behavior.

## Introduction

Rhythm generation is an essential function of the nervous system. Rhythmic neuronal firing is vital to nutrient intake (Marder & Calabrese, 1996; Spencer et al., 2018), waste expulsion (Bywater et al., 1989), breathing (Feldman, 1986), and locomotion (Dougherty & Ha, 2019). Aberrant neuronal rhythms also arise in pathologies, such as in neuropathic pain (Amir et al., 1999) and Parkinson’s Disease (Oswal et al., 2013). Within rhythmic motor behaviors including mastication, respiration, and locomotion, neurons employ rhythmogenic network and intrinsic properties to encode the timing of motor output (Harris-Warrick, 2010). In the spinal cord, the locomotor central pattern generator (CPG) includes dedicated neurons for rhythmic locomotor output (McCrea & Rybak, 2008). Neonatal rodents have been used in many studies of neurons involved in locomotor circuitry due to transgenic neuronal targeting and the ability to evoke fictive locomotor activity in reduced neonatal preparations (Cazalets et al., 1996; Cowley & Schmidt, 1997; Dougherty et al., 2013; Kjaerulff & Kiehn, 1996; Ziskind-Conhaim et al., 2010). While capable of producing locomotor-like behavior, the neonatal spinal cord is immature in its cellular and synaptic properties (Borowska et al., 2015; Chang et al., 1999; Ha & Dougherty, 2018; Sharples & Miles, 2021). However, despite developmental changes within the spinal locomotor CPG, locomotor rhythm generating neurons are thought to remain rhythmogenic throughout the life of the animal. Whether significant changes developmentally occur in the intrinsic rhythmogenic properties of locomotor CPG neurons is beginning to be explored (Husch et al., 2015).

Spinal Shox2 interneurons (INs) are a putative rhythm-generating population in the mouse spinal cord (Dougherty, 2023) and are positioned to be rhythm-generating in that they are ventromedially located, excitatory, and form mutual connections (Brownstone & Wilson, 2008). Furthermore, silencing these neurons decreases the frequency of locomotor-like activity in the isolated neonatal spinal cord (Dougherty et al., 2013). Previous work has shown that neonatal Shox2 INs are highly interconnected by gap junctional coupling, with decreasing prevalence as the mouse ages (Ha & Dougherty, 2018). Rhythmicity is evident in neonatal Shox2 INs during fictive locomotion (Dougherty et al., 2013) but in the adult spinal slice, where network connections are limited to those that are local within the slice, a small subset of Shox2 INs display spontaneous oscillations (Garcia-Ramirez et al., 2021). These findings indicate that Shox2 INs are rhythmogenic even within a limited network and potentially employ intrinsic rhythmogenic properties which change with age. Although lumbar spinal Shox2 INs are a heterogeneous population (Garcia-Ramirez et al., 2022) and are not the only locomotor-related rhythmogenic neuron type (Chalif et al., 2022; Ziskind-Conhaim et al., 2010), we use them in the present study as a representative neuronal population for rhythmogenic spinal CPG neurons.

Rhythmicity of CPG neurons, like in many other rhythmogenic neuron types across the nervous system, is maintained and shaped, at least in part, by potentially rhythmogenic ionic currents which shape and maintain it (Harris-Warrick, 2010). These include slowly inactivating inward currents which maintain the sustained depolarization needed for burst firing, such as persistent inward currents (PICs), hyperpolarization-activated current (I_H_), and T-type calcium current (I_T_) (Harris Warrick, 2002; Harris-Warrick, 2010; Verneuil et al., 2020). To balance this, outward or hyperpolarizing currents also shape rhythmic firing, such as slow afterhyperpolarization (sAHP), and M- (I_M_) and A-type (I_A_) potassium currents (Harris Warrick, 2002; Harris-Warrick, 2010; Verneuil et al., 2020). These putatively rhythmogenic currents may be employed in a myriad of combinations across different modeled biological contexts (Alonso & Marder, 2020). Additionally, each ionic current is mediated within individual neurons by a suite of voltage-sensitive ion channels which we are beginning to understand using genetic and histological tools. Moreover, the rhythmic firing in neurons is determined by the combination of passive and active cellular properties. Within these complexities of rhythmogenic intrinsic properties may lie future therapeutic targets for the modulation or restoration of rhythm in the spinal locomotor CPG following injury (Brocard et al., 2016). Accordingly, it is crucial to understand rhythmogenic spinal CPG neurons in adult animals, where cellular properties and connections have reached maturity.

In this study, we sought to determine which active ionic currents are displayed by Shox2 INs in mice and how they change with postnatal development. Using whole-cell patch clamp electrophysiology, we found that all examined putatively-rhythmogenic currents are displayed in small or large subsets of Shox2 INs. Strong PICs which persist onto the descending phase of a symmetrical voltage ramp and sAHPs are largely absent in Shox2 INs from neonatal mice. We found that I_M_, like PICs, increases in magnitude with age. Further, subsets of Shox2 INs across postnatal development express channels which are known to contribute to PICs, I_M_, and sAHP. Since it is the combination of passive and active cellular properties that determine the interactions resulting in intrinsic rhythm, we performed unbiased hierarchical clustering of all measured electrophysiological parameters. This revealed electrophysiological subtypes of Shox2 INs that were age-dependent. The subtypes found in older mice were identified by increased diversity and strong active current expression, suggesting that these properties are enhanced with postnatal age in putative rhythm generating neurons.

## Materials and Methods

### Mouse Lines

All animal experiments were performed using the following transgenic mouse lines: *Shox2::Cre* (Dougherty et al., 2013); *R26-lsl-tdTomato* (Ai9 from The Jackson Laboratory, #007909, Madisen et al., 2010) and *Shox2::Cre; R26-lsl-tdTomato;Vsx2-eGFP* (Vsx2-EGFP from Mutant Mouse Regional Resource Center, #011391-UCD, (Gong et al., 2003). Neonatal (P2-5, 7), juvenile (P14-21), and adult (P60-138) transgenic mice were used for this study. All experimental procedures followed National Institutes of Health guidelines and were approved by the Institutional Animal Care and Use Committee at Drexel University.

### Slice Preparation

To access lumbar spinal Shox2 INs in the spinal slice, juvenile and adult mice were first anesthetized with ketamine (150mg/kg) and xylazine (15mg/kg). Following decapitation and evisceration, spinal cords were removed from all mice in ice-cold dissecting solution. The dissecting solution used for neonatal mice (P2-5) contained the following (in mM): 111 NaCl, 3 KCl, 11 glucose, 25 NaHCO_3_, 3.7 MgSO_4_, 1.1 KH_2_PO_4_, and 0.25 CaCl_2_. For juvenile and adult mice, the dissection solution contained (in mM): 222 glycerol, 3 KCl, 11 glucose, 25 NaHCO_3_, 1.3 MgSO_4_, 1.1 KH_2_PO_4_, and 2.5 CaCl_2_. The lumbar spinal cord (L1-5) was sectioned transversely (300µm) in dissection solution using a vibrating microtome (Leica Microsystems). Slices were immediately transferred to recording artificial cerebrospinal fluid (ACSF) containing the following (in mM): 111 NaCl, 3 KCl, 11 glucose, 25 NaHCO_3_, 1.3 MgSO_4_, 1.1 KH_2_PO_4_, and 2.5 CaCl_2_. Slices from a subset of juvenile and all adult mice were incubated at 34-37°C for 20-30 minutes and then rested at room temperature for 1 hour before recording. Dissecting and recording solutions were continuously aerated with 95%/5% O_2_/CO_2_.

### Electrophysiological Recordings

Fluorescently labeled tdTomato^+^ Shox2 INs were visualized with a 63X objective lens on a BX51WI scope (Olympus) using LED illumination (Lumen Dynamics X-Cite) and targeted for whole cell patch clamp recordings. Electrodes were pulled to tip resistances of 5–12 MΩ using a multi-stage puller (Sutter Instruments) and were filled with intracellular solution which contained (in mM): 128 K-gluconate, 10 HEPES, 0.0001 CaCl_2_, 1 glucose, 4 NaCl, 5 ATP, and 0.3 GTP. In experiments using riluzole or nimodipine, a cesium-based intracellular solution was used, containing (in mM): 130 CsMeSO_3_, 10 HEPES, 0.5 EGTA, 2 MgCl_2_, 2 ATP, 5 CsCl. In some experiments, biocytin (2 mg/mL, Sigma) was included in the patch electrode. All recordings were performed at room temperature. Data were collected with a Multiclamp 700B amplifier (Molecular Devices) and Clampex software (pClamp9, Molecular Devices). Signals were digitized at 20kHz and filtered at 6kHz. Measurements were manually calculated from recording traces in Clampfit 11 (Clampex, Molecular Devices).

### Electrophysiological Data Collection and Analysis

Resting membrane potential was recorded shortly after gaining whole-cell access. Recordings in current-clamp were taken using current steps while holding the cell near -65mV via injecting bias current. Input resistance was calculated from the current/voltage slope in response to hyperpolarizing current steps in which no voltage-gated current activation was present. Time constant was measured as the duration of time to reach 63% of the maximum membrane voltage change in response to a hyperpolarizing current step. Input capacitance was calculated by dividing input resistance by time constant. Leak conductance was calculated as the inverse of the input resistance. Action potential properties (threshold, rheobase, sAHP duration, and sAHP amplitude) were measured from the first spike elicited from the depolarizing current step at rheobase. Voltage-sensitive currents were isolated in voltage-clamp mode. To record PICs, cells were held at -80mV and subjected to three sequential symmetrical voltage ramps in which the cells were brought to -10mV in 2.5 seconds, and back down to -80mV in 2.5 seconds (Dai & Jordan, 2010). Measurements including PIC onset, area of the current, and maximum amplitudes were taken from the first voltage ramp. To isolate and measure I_H_, cells were subjected to a 500ms hyperpolarizing step from -60mV to -140mV. Occasionally, a cell would not return to steady state within the 500ms duration, so 1 second steps were performed in addition. To record and measure I_T_ and I_A_, cells were subjected to two sequential 250ms depolarizing steps to -30mV, the first was from -100mV to deinactivate the relevant channels and the second from -60mV. Measurements were taken from the voltage step that the current was maximally activated by. The tail relaxation curve of I_M_ was measured from recordings of cells hyperpolarized to -60mV from an initial holding of -10mV (Brown & Adams, 1980). Current magnitudes are reported as current density, where the maximum amplitude of the current is normalized by the cell’s input capacitance.

### Pharmacology

The following pharmacological agents were used in this study: 10μM riluzole (Sigma R116), 20μM nimodipine (Sigma N149), 10μM XE991 (Sigma X2254), and 100nM apamin (Sigma A1289). All were dissolved in recording ACSF except for riluzole, which was dissolved in DMSO and subsequently added to recording ACSF. The final concentration of DMSO was 0.05%.

### Immunohistochemistry and RNAscope *in situ* Hybridization

*Shox2::Cre;lsl-tdTomato* mice at P14 and P64 were anesthetized with ketamine (150mg/kg) and xylazine (15mg/kg) and perfused transcardially with 0.1M PBS, followed by 4% PFA in PBS. *Shox2::Cre;lsl-tdTomato* mice at P2 and P7 were decapitated and eviscerated in PBS. Spinal cords were harvested from each animal and fixed overnight in 4% PFA solution at 4°C. Fixed spinal cords were subsequently maintained in 30% sucrose in PBS for at least 48 hours. Tissue was then embedded in OCT compound (Thermo Fisher Scientific) over dry ice and stored at -80°C. Lumbar spinal cords were sectioned (20 µm) transversely on a cryostat (Microm HM 505 E) and directly mounted onto coated (immunohistochemistry) or charged (RNAscope) slides. Slides were washed in PBS before being used for immunohistochemistry or RNAscope.

For immunohistochemistry, slides were first blocked in a PBS solution containing 5% goat serum, 1% bovine serum albumin, 0.2% Triton X-100, and 0.1% fish gelatin. Slides were incubated in one of the following primary antibodies overnight: rabbit anti-KCNQ2 (1:1000, ThermoFisher PA1-929), rabbit anti-KCNQ3 (1:400, Alomone APC-051), rabbit anti-SK2 (1:50 Alomone APC-028), or rabbit anti-SK3 (1:50, Alomone APC-025). Slides were then incubated for 2 hours in goat anti-rabbit 647 secondary antibody (1:400, Invitrogen A-21245). All immunohistochemistry steps were performed at room temperature.

RNAscope was performed according to manufacturer’s protocols (Wang et al., 2012). ACDBio probes used include tdTomato (317041), SCN8a (434191-C2), and CACNA1d (502591-C2). All slides from both immunohistochemistry and RNAscope were coverslipped using Fluoromount-G with DAPI (Invitrogen 00-4959-52). Images were acquired as sequential z stacks of 20x tile-scans on a Leica Microsystems SP8 confocal microscope. Cell counting was performed manually in ImageJ using the multipoint tool. All Shox2 INs in each examined slice were counted. Slices (n=7-12 per animal), at least 200 µm apart, from lumbar segments of the spinal cord were quantified.

### Agglomerative Hierarchical Clustering

Pearson’s linear correlation test was conducted to exclude highly correlated electrophysiological parameters (*p*<0.0001) from the 18 which are detailed above. Hierarchical cluster analyses were performed on the 11 electrophysiological parameters that were determined to be not highly correlated among each other. These 11 parameters included resting membrane potential, time constant, input capacitance, action potential threshold, sAHP amplitude, PIC area, PIC onset voltage, PIC with descending density, I_H_ area, I_T_ density, and I_M_ density. Hierarchical clusters were determined by the correlation distance between pairs of observations and consideration of an average for the distance between clusters. This combination was used because it resulted in the highest cophenetic correlation coefficient (0.3046). The number of clusters was determined considering the cutoff below the maximum inconsistency coefficient for each link. Data was initially standardized to a mean of 0 and standard deviation of 1 to be able to compare variables with different units in MATLAB using customized scripts.

### Statistical Analyses

Statistical tests and post hoc analyses were performed with GraphPad Prism (GraphPad Software). All results are presented as mean ± SD, unless otherwise stated. Statistical significance was set at p<0.05. The distribution of the data was determined by Shapiro–Wilk normality test. In the case of normally distributed data, we performed paired t-tests to compare baseline and drug conditions or one-way ANOVAs with Tukey post hoc tests to compare 3 age groups. In the cases where data were not normally distributed, we performed Wilcoxon matched pairs signed ranks tests to compare baseline and drug conditions and Kruskal-Wallis with Dunn’s multiple comparisons post hoc tests to compare 3 age groups. Comparisons between age groups of percentages for conductance presence, RNA presence, or protein presence were performed with chi-square test.

## Results

### Intrinsic passive and active properties of Shox2 INs during development

Spinal Shox2 INs are part of the locomotor CPG that generates rhythmic and patterned motor output across the lifespan (Dougherty et al., 2013). Although mice begin performing regular weight-bearing steps by P9 (Jiang et al., 1999a), mainly neonatal mice have been used in previous studies of spinal locomotor circuitry (i.e. Anderson et al., 2012; Dougherty et al., 2013; Hinckley et al., 2010). The general neuronal types and connectivity structure in the neonatal spinal cord are hypothesized to be present and similar in function to those in adult mice (Crone et al., 2008, 2009; Rybak et al., 2015; Danner et al., 2017, Talpalar et al., 2013). However, there are some notable differences. Given that strong gap junctional coupling in neonatal locomotor circuits decreases by adulthood (Ha & Dougherty, 2018), we hypothesized that some intrinsic rhythmogenic properties in Shox2 INs increase in magnitude and/or prevalence with age. Such a developmental change may compensate for reduced gap junctional coupling and maintain locomotor rhythm in adults. To compare properties of these neurons at neonatal, adult, and an intermediate juvenile stage, we first examined the basic electrophysiological properties of Shox2 INs across postnatal maturation. We recorded passive and active properties using whole-cell patch clamp in lumbar spinal slices targeting 35 neonatal (P2-5, mean age 3.2 ± 0.980), 33 juvenile (P14-21, mean age 17.2 ± 1.61), and 36 adult (P60+, mean age 96.2 ± 28.52) fluorescently labeled Shox2 INs (Table 1). We found that neonatal cells have a more depolarized action potential threshold voltage (-30.5 ± 4.3 mV, n=35) compared to adult (-33.2 ± 4.7 mV, n=36) Shox2 INs (one-way ANOVA with Tukey’s *post hoc* test, p=0.0279). Neonatal Shox2 INs have a longer time constant (53.0 ± 29.5 ms, n=35) compared to juvenile (33.4 ± 12.4 ms, n=33; Kruskal-Wallis with Dunn’s *post hoc* test, p=0.0021) and adult (31.2 ± 17.0 ms, n=36; Kruskal-Wallis with Dunn’s *post hoc* tes,t p<0.0001) Shox2 INs. We also report a higher input capacitance in neonatal Shox2 INs (54.3 ± 17.3 pF, n=35) compared to juvenile (47.1 ± 25.3 pF, n=32; one-way ANOVA with Tukey’s *post hoc* test, p=0.0149) and adult (43.1 ± 16.8 pF, n=36; one-way ANOVA with Tukey’s *post hoc* test, p=0.0104) Shox2 INs, which is consistent with the findings reported for lumbar spinal V2a INs (Husch et al., 2015). We found no significant differences between the age groups in resting membrane potential, input resistance, rheobase, and leak conductance. These findings indicate relatively modest differences in active and passive electrophysiological properties in lumbar spinal Shox2 INs across postnatal age which are consistent with previously reported developmental changes in lumbar spinal INs (Borowska et al., 2015, Husch et al., 2015).

**Table 1.**
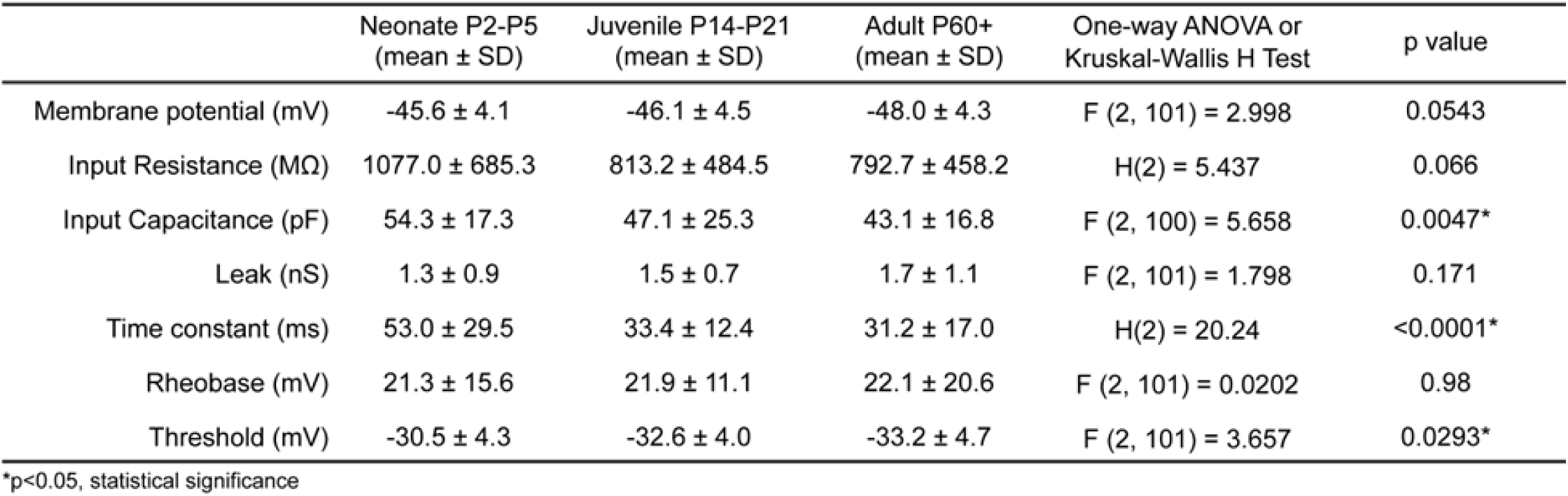
Comparison of intrinsic passive and active properties in Shox2 INs from three age groups.

### The hyperpolarization-activated, T-type calcium, and A-type potassium currents are found in subsets of Shox2 INs across postnatal age

To reveal the extent to which Shox2 INs display voltage-sensitive rhythmogenic properties across postnatal development, we recorded voltage-sensitive conductances that have been previously shown to shape bursting and firing properties in other neuron types (Harris-Warrick, 2010). This included I_H_ (Fig. 1*A*), which was present in approximately half of the recorded neonate (n=20/53, 57%) and adult (n=18/36, 50%) Shox2 INs, and a slightly larger subset of juvenile (n=21/32, 66%) Shox2 INs (Fig. 1*B*). The current density of I_H_ did not change across postnatal age groups (Fig. 1*C*). Both I_T_ (Fig. 1*D, E*), and I_A_ (Fig. 1*G, H*) were present in ≤17% of Shox2 INs across all three age groups. The current density of I_T_ does increase from neonate (0.44 ± 0.23 pA/pF, n=6) to adult (1.13 ± 0.50 pA/pF, n=10; one-way ANOVA with Tukey’s *post hoc* test, p=0.0179) Shox2 INs, despite being a rarely displayed property in these neurons (Fig. 1*F*). The current density of I_A_ did not change across postnatal age groups (Fig. 1*I*). These findings reveal that all three tested conductances occur in subsets of Shox2 INs from all age groups.

**Figure 1.**
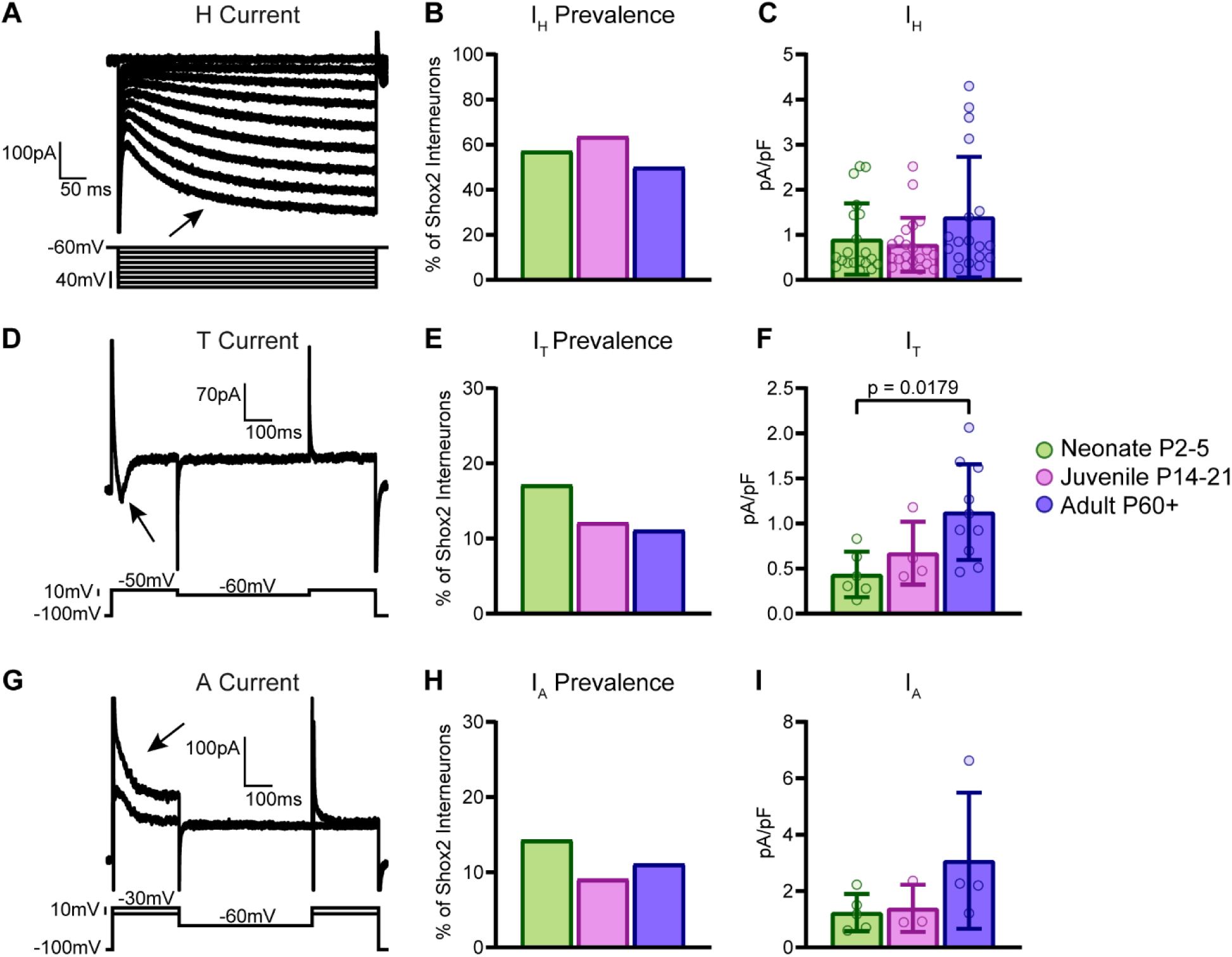
I_H,_ I_T,_ and I_A_ are found in subsets of Shox2 INs. (A) Inward current responses to hyperpolarizing voltage steps from -60mV. Arrow indicates the presence of I_H_. (B) 50-66% of Shox2 INs display I_H_. (C) I_H_ current densities are relatively consistent through postnatal development. Error bars indicate SD. (D) Inward current response to depolarizing voltage step from -100mV, and not from -60mV. Arrow indicates the presence of I_T_.(E) <18% of Shox2 INs display I_T_. (F) I_T_ current density increases from neonate to adult. One-way ANOVA with Tukey *post-hoc* test. Error bars indicate SD. (G) Outward current response to depolarizing voltage step from -100mV, and not from -60mV. Arrow indicates the presence of I_A_. (H) <15% of Shox2 INs display I_A_. (I) I_A_ current densities do not change significantly with age. Error bars indicate SD.

### Shox2 INs display PICs which increase in magnitude with age, and express RNA for PIC-related ion channels

To record the presence of persistent depolarizing conductances, we used voltage ramps to isolate PICs in Shox2 INs (Fig. 2*A, B*). We found that most neonatal (n=27/35, 77%), juvenile (n=29/32, 91%), and adult (n=31/36, 86%) Shox2 INs display a persistent inward current (PIC) in response to slow current ramp (Fig. 2*C*). In addition to PICs being present on the ascending phase of the voltage ramp, we found that a small subset of Shox2 INs display PICs on both the ascending and descending phases of the ramp (Fig. 2*D*). In these cases, the current onset occurs on the ascending phase of the ramp, while the offset occurs on the descending phase (Fig. 2*B*). Of note, a small subset of PICs in juvenile (n=6/32, 19%) and adult (n=6/36, 17%) display PICs on the descending ramp, which is a feature largely missing from neonatal (n=1/35, 3%) Shox2 INs (Fig. 2*D*). This may be due to either the strength of the PIC that carries over the duration of the voltage ramp, the time constant of the slow inactivation of PICs, or a shifted voltage dependence of deinactivation in the subset of Shox2 INs displaying the descending PICs.

**Figure 2.**
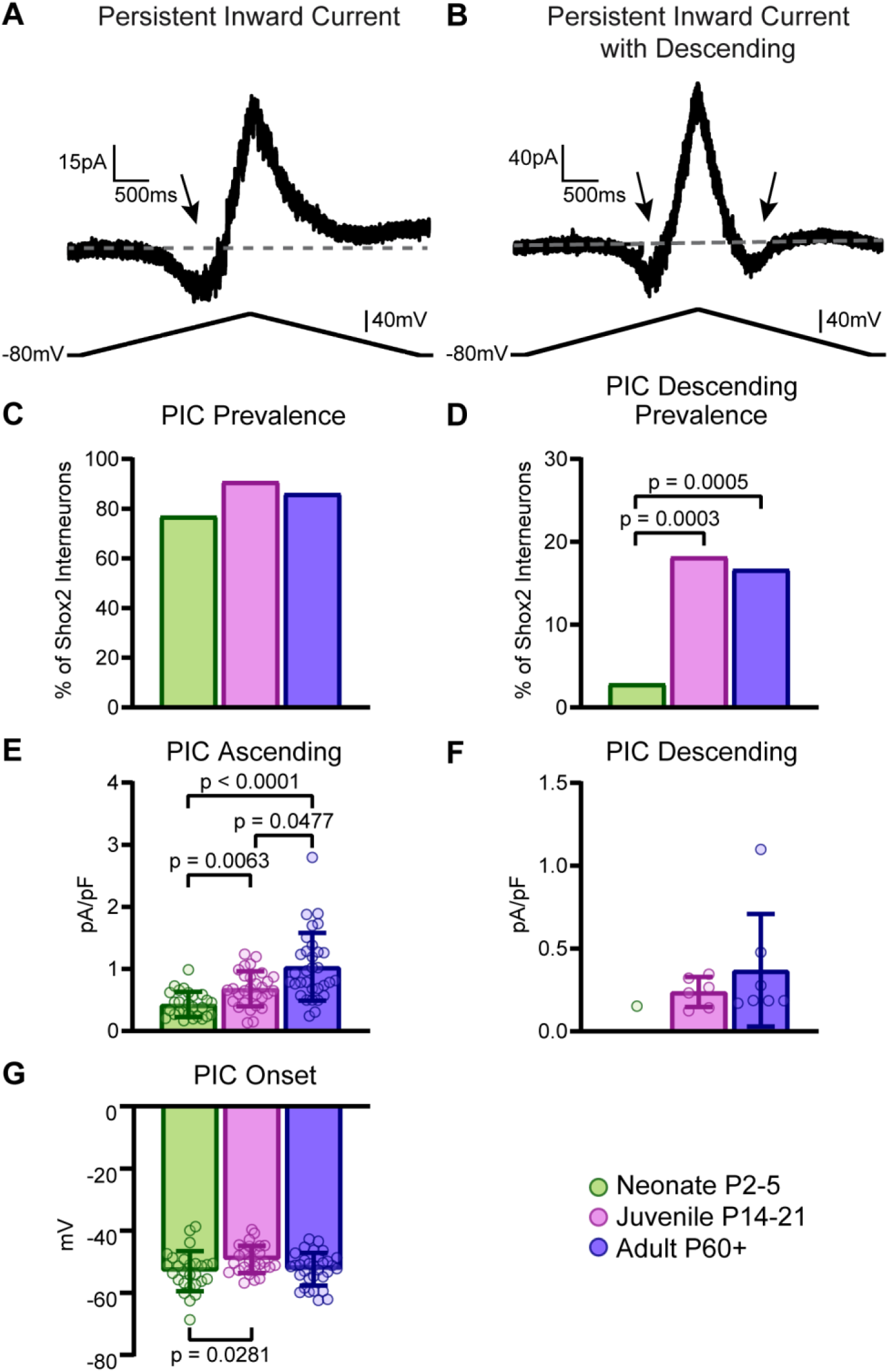
Subsets of Shox2 INs display PICs which increase in magnitude with age. (A, B) Inward current responses to slow voltage ramps from -80mV to -10mV back to -80mV. Arrows indicate the presence of PICs, some of which are only present in the first part of the ramp (A) and others persist in the descending phase of the ramp (B). (C) PICs are present in most (<75%) Shox2 INs from all age groups. (D) PICs with descending are less prevalent (<20%) in juvenile and adult Shox2 INs and are even less prevalent in neonate Shox2 INs (3%) compared to juvenile (Chi-square, DF 2, p=0.0003) or adult (Chi-square, DF 2, p=0.0005). (E) PICs increase in current density from neonate (green) to juvenile (pink) to adult (purple) Shox2 INs. (F) Current densities of the relatively few Shox2 INs which display PIC with descending. Kruskal-Wallis with Dunn’s *post-hoc* test. Error bars indicate SD. (G) Mean onset of PICs in response to the ascending voltage ramp is more depolarized in juvenile compared to neonateShox2 INs. One-way ANOVA with Tukey *post-hoc* test. Error bars indicate SD.

As PICs can be mediated by both sodium and calcium components, we further examined the voltage properties and pharmacological sensitivities of PICs in Shox2 INs across postnatal development. The current density of PICs on the ascending phase of the voltage ramp increases from neonate (0.42 ± 0.20 pA/pF, n=27) to juvenile (0.68 ± 0.28 pA/pF, n=29; Kruskal-Wallis test with Dunn’s *post hoc* test, p=0.0063) and juvenile to adult (1.03 ± .54 pA/pF, n=31; Kruskal-Wallis test with Dunn’s *post hoc* test, p=0.0477, Fig. 2*E*). As PICs persisting onto the descending phase of the voltage ramp were rarely present, particularly in neonates (Fig. 2*D*), we were unable to test for a statistical change in its magnitude with development (Fig. 2*F*). We found that PIC onset is more depolarized in juvenile (-49 ± 4.5 mV, n=29) compared to neonatal Shox2 INs (-53 ± 6.3 mV, n=27; one-way ANOVA with Tukey’s *post hoc* test, p=0.0281), which are similar to the adult Shox2 INs (-52 ± 5.2 mV, n=31, Fig. 2*G*). These data demonstrate that there are developmental magnitude increases in PICs in Shox2 INs. We then asked whether PICs in adult Shox2 INs consist of persistent sodium (I_NaP_) and L-type calcium (I_Ca,L_) components. We applied 10μM riluzole (Fig. 3*A*) or 20μM nimodipine (Fig. 3*B*) to block these currents respectively in adult Shox2 INs, using an intracellular solution containing cesium to increase space clamp. With the inclusion of cesium, all (n=20/20) Shox2 INs displayed PICs on the descending phase of the ramp. Riluzole (n=12) and nimodipine (n=8) application resulted in a mean decrease of 34% and 14% in current amplitude, respectively (Fig. 3*C, D*). However, only riluzole application resulted in a decreased PIC on the descending phase of the ramp (Fig. 3*C*, right).

**Figure 3.**
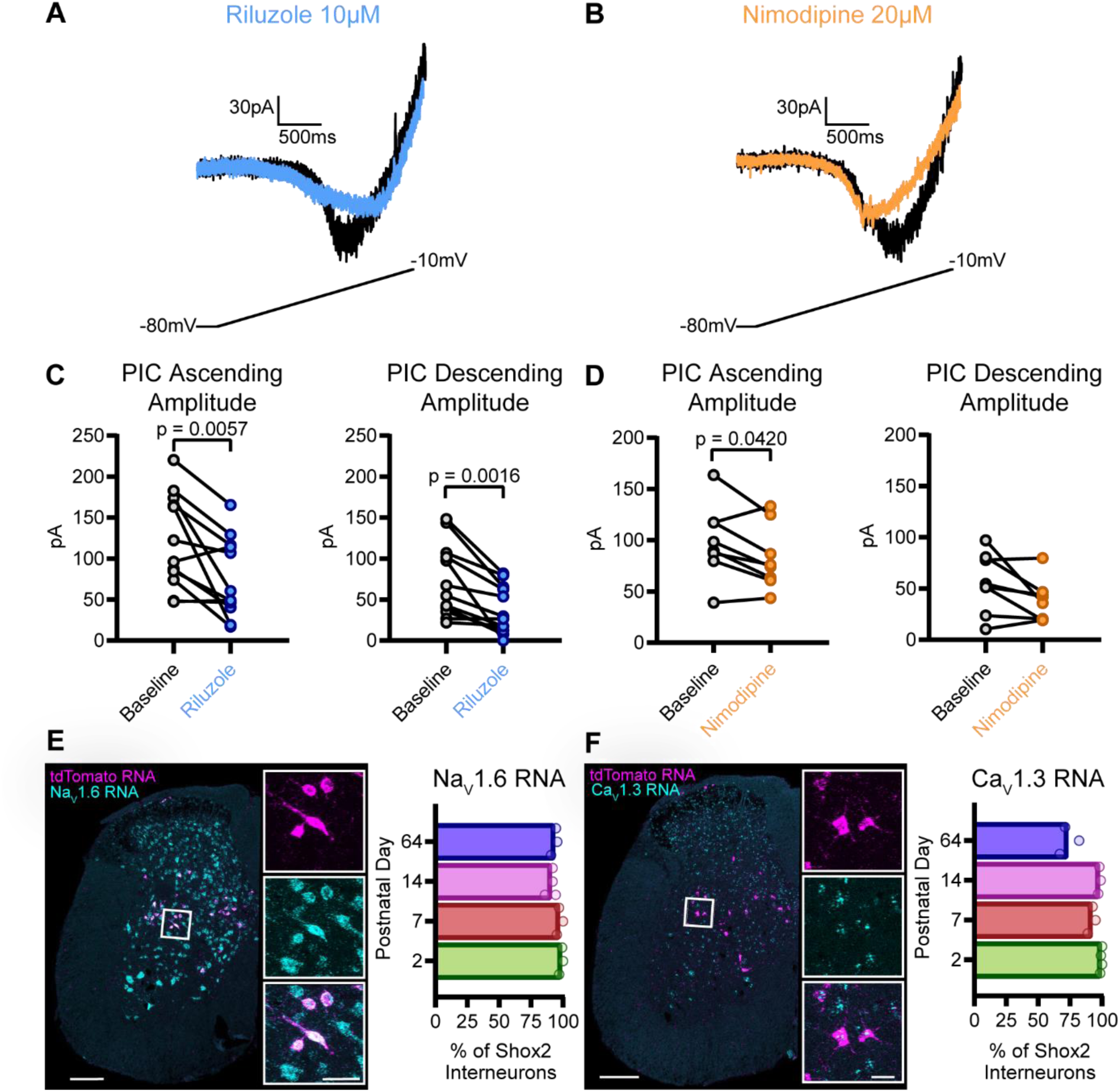
Subsets of Shox2 INs display PICs which have both sodium and calcium components. (A) Representative examples of PICs at baseline (black) and following riluzole application (blue). (B) Representative examples of PICs at baseline (black) and following nimodipine application (orange). (C) Riluzole decreases the amplitudes of PICs measured in both the ascending and descending phases of the ramp, paired t-test. (D) Nimodipine decreases the amplitudes of PICs but not the descending phase of PICs, paired t-test. (E) Representative images of RNA for Na_V_1.6 (cyan) and tdTomato (magenta) in the lumbar spinal slice. >85% of tdTomato RNA-containing cells also contain NA_V_1.6 RNA across all four ages examined. (F) Representative images of RNA for Ca_V_1.3 (cyan) and tdTomato (magenta) in the lumbar spinal slice. Lumbar spinal sections from P64 have less tdTomato RNA-containing cells that also contain Ca_V_1.3 RNA than the other three ages examined. Scale bars for spinal sections=200µm, insets=25µm.

Having observed both sodium and calcium components of PICs, we next investigated whether lumbar spinal Shox2 INs display the RNA transcripts correlated to these conductances. We examined four postnatal ages: 2, 7, 14, and 64 days. We began with Na_V_1.6, a voltage-gated sodium channel responsible for mediating I_NaP_ in other neuron populations (Katz et al., 2018; Sharples & Miles, 2021). Using immunohistochemistry, we were unable to visualize Na_V_1.6 expression colocalized to tdTomato positive Shox2 INs which is likely due to predominant expression patterns in the axon initial segments of neurons (Drouillas et al., 2023). Visualization of Shox2 IN axon initial segments is difficult due to their mixed orientations, and a lack of tdTomato expression in cellular compartments beyond the soma. Therefore, we performed RNAscope *in situ* hybridization to visualize RNA within lumbar spinal Shox2 IN cell bodies at four ages. With this method, we found that Na_V_1.6 RNA is highly prevalent in Shox2 INs at all ages examined (Fig. 3*E*). We next targeted Ca_V_1.3, a voltage-gated calcium channel which mediates I_Ca,L_ (Jiang, et al., 1999b). Ca_V_1.3 RNA is present in a large proportion (<67%) of examined lumbar spinal Shox2 INs at 2, 7, 14, and 64 postnatal days (Fig. 3*F*).

Together, these data show that Shox2 INs display sodium- and calcium-mediated PICs which increase in amplitude with age, as well as RNA transcripts for channels which can mediate both I_NaP_ and I_Ca,L_ components of PICs.

### Shox2 INs display M currents which increase in magnitude with age and voltage-gated potassium channels

I_M_ (Fig. 4*A*) is the most prevalent current in Shox2 INs among all examined ionic currents and across all three age groups (Fig. 4*B*). We found that most neonatal (n=31/35, 87%), juvenile (n=28/32, 88%), and adult (n=31/36, 86%) Shox2 INs display the tail relaxation curve from I_M_ (Fig. 4*B*). We hypothesized that this property increases in magnitude with age. We found that I_M_ does increase in magnitude between neonate (1.17 ± 0.44 pA/pF, n=31) and adult (2.26 ± 1.7 pA/pF, n=31; Kruskal-Wallis test with Dunn’s *post hoc* test, p=0.0173, Fig. 4*C*) Shox2 INs, with juvenile I_M_ magnitude residing between (1.78 ± 1.08, n=28). I_M_ is mediated by heteromers of two voltage-gated potassium channel subunits: K_V_7.2 and K_V_7.3 (Wang et al., 1998). We blocked K_V_7.2/3 channels by applying XE991 (Greene et al., 2017) and measured a mean 51% decrease in I_M_ amplitude, confirming that this current is mediated by K_V_7.2/3 channels in Shox2 INs (mean age 4.25 ± 3.03, Fig. 4*D*). We visualized these channels in lumbar spinal Shox2 INs across postnatal development using immunohistochemistry and found that Shox2 INs express both channel subunits at all ages examined (Fig. 4*E*, *F*). These data show that K_V_7.2/3-mediated I_M_ increases in magnitude across postnatal development in Shox2 INs, and that Shox2 INs express K_V_7.2/3 channels from neonatal ages through adulthood.

**Figure 4.**
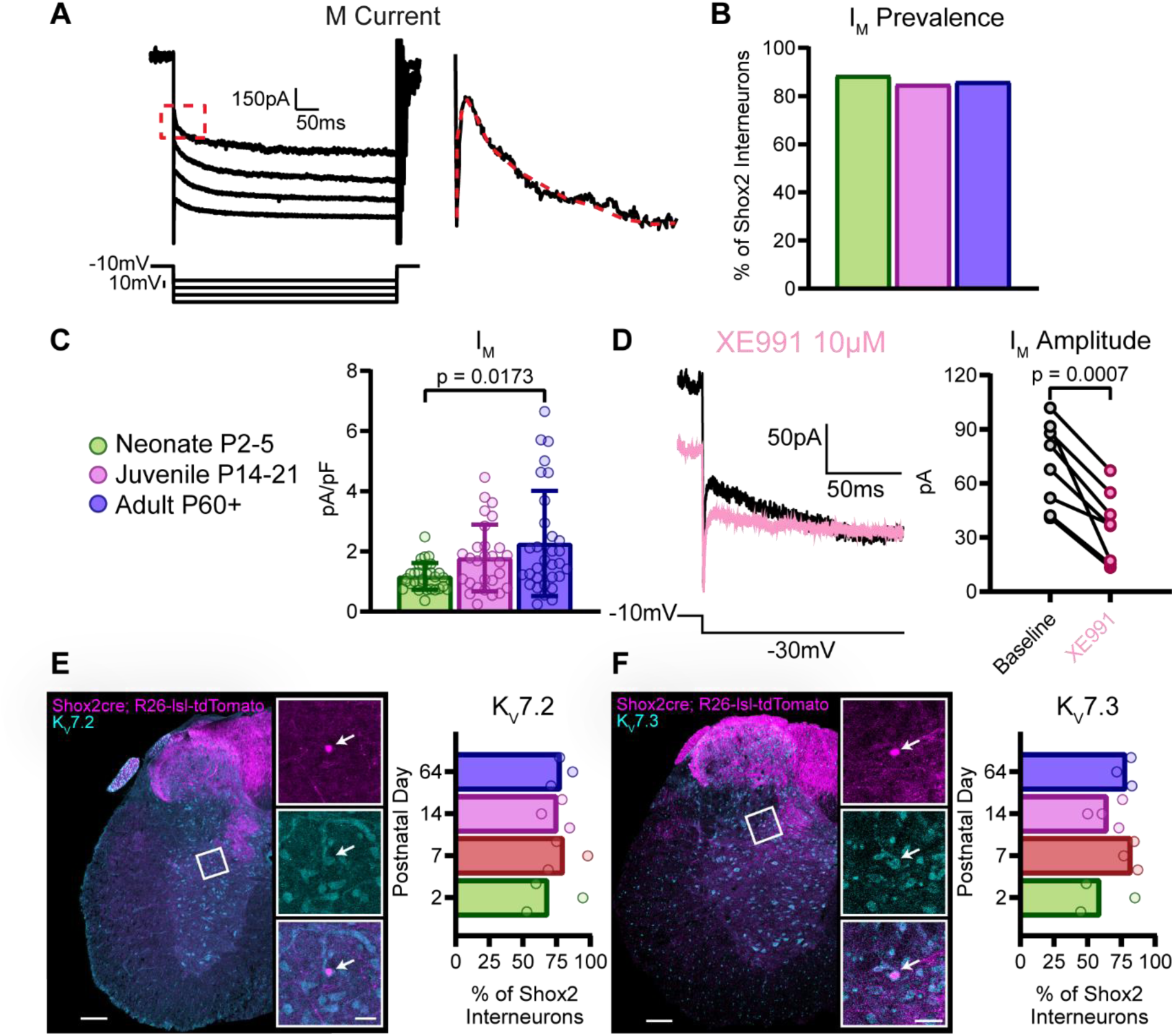
Subsets of Shox2 INs display M currents which increase in magnitude with age and express K_V_7.2/3 subunits. (A) Outward current responses to hyperpolarizing voltage steps from -10mV. Red dotted box and line highlight tail relaxation curve of of I_M_. (B) I_M_ is the most prevalent property (≥85%) in Shox2 INs across all age groups among those examined in this study. (C) I_M_ current density increases from neonate (green) to adult (purple). Kruskal-Wallis with Dunn’s *post-hoc* test. Error bars indicate SD. (D) Representative I_M_ tails from baseline (black) and XE991 (pink) conditions. XE991 decreases the amplitude of I_M_ tails. Paired t-test. (E, F) Representative images and quantification of (E) K_V_7.2 and (F) K_V_7.3 expression using immunohistochemistry. Channel expression is cyan and tdTomato expression signifying Shox2 is magenta. ≥50% of tdTomato^+^ neurons express K_V_7.2 or K_V_7.3 across all ages examined. Scale bars for spinal sections=200µm, insets=25µm.

### Juvenile and adult Shox2 INs display sAHP and express calcium-dependent potassium channels

We observed biphasic AHP, indicative of sAHP, in current-clamp (Fig. 5*A*) in a subset of juvenile (n=18/32, 56%) and adult (n=20/36, 56%) Shox2 INs, and rarely in neonatal INs (n=1/35, 3%, Fig. 5*B*). The amplitude (Fig. 5*C*) and duration (Fig. 5*D*) of the sAHP was similar between juvenile and adult Shox2 INs. We also found that the sAHP in adult Shox2 INs is sensitive to 100nM apamin (Fig. 5*E*), a blocker for SK2 and SK3 calcium-dependent potassium channels (Köhler et al., 1996), as it completely eliminated the sAHP in all but one tested Shox2 IN (n=9/10, Fig. 5*F, G*). As sAHP increases in prevalence at juvenile and adult stages, we hypothesized that the channels mediating it would also increase in prevalence with age in Shox2 INs. Motor neurons have been shown to display an apamin-sensitive sAHP mediated by the calcium-dependent potassium channels SK2 and SK3 (Deardorff et al., 2013), so we examined these channels in Shox2 INs using immunohistochemistry. We found that SK2 (Fig. 5*H*) and SK3 (Fig. 5*I*) channels are rarely present in examined Shox2 INs from the youngest ages studied here, P2 (2%-17%) and P7 (5%-16%). However, 30% of juvenile (P14) and adult (P64) Shox2 INs express SK2 channels and 31% express SK3 channels (Fig. 5*H, I*). These SK channels have robust expression in other spinal neuron types as well. Taken together, we show that sAHP magnitude and SK2/3 channel expression are similarly increased in Shox2 INs at juvenile and adult time points.

**Figure 5.**
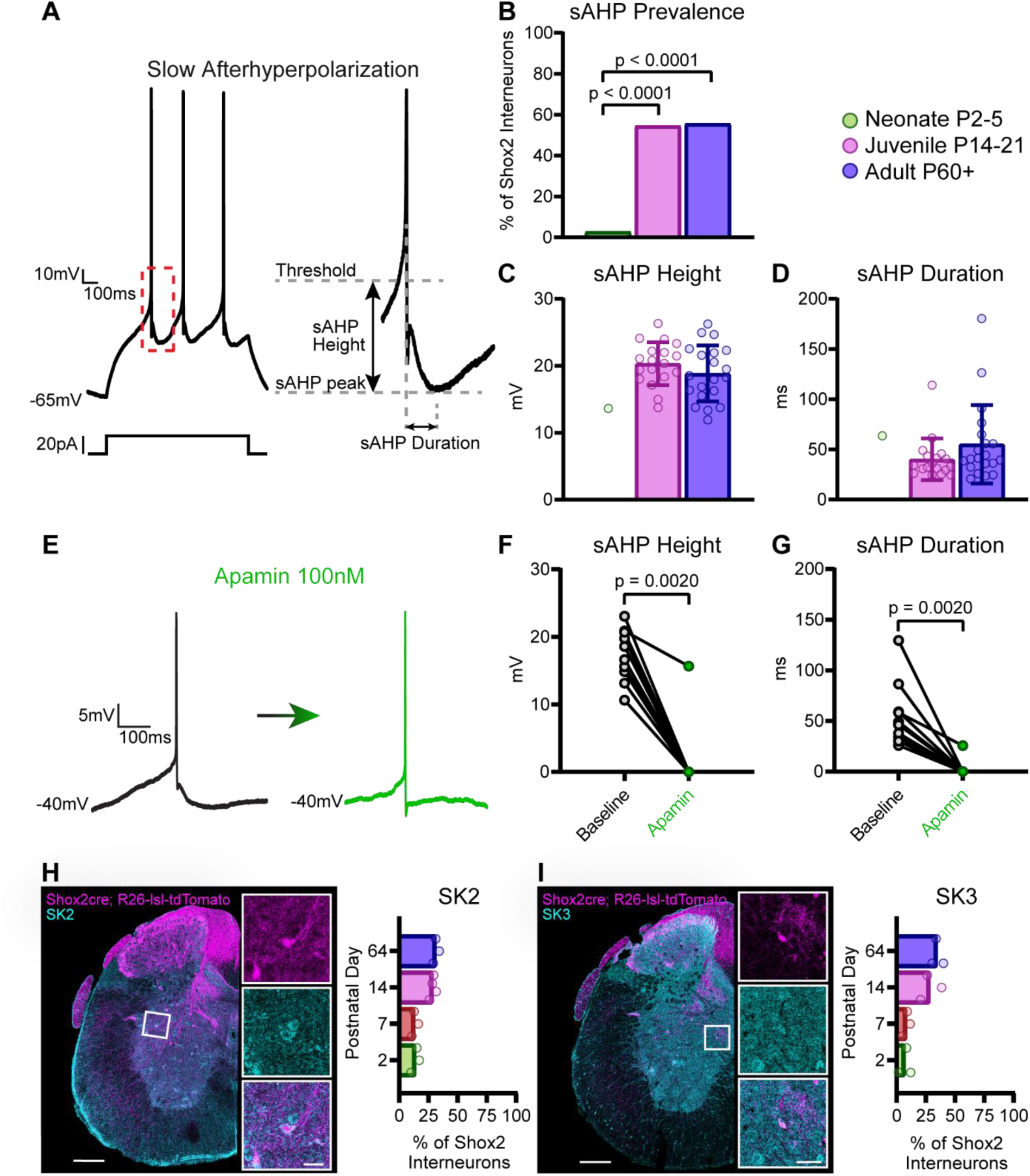
Subsets of juvenile and adult Shox2 INs display sAHP and express SK2 and SK3. (A) Representative example of action potentials with sAHP in response to depolarizing current step. Red dotted box indicates inset highlighting the sAHP, with height and duration measured as shown. (B) sAHP is rare in neonate Shox2 INs compared to juvenile (Chi-square, DF 2, p<0.0001) or adult (Chi-square, DF 2, p<0.0001). (C) Comparison of sAHP height and (D) duration, which is consistent between juvenile and adult stages. Error bars indicate SD. (E) Example of an action potential at baseline with a sAHP (black), and following application of apamin, which blocks the sAHP (green). (F) Apamin decreases both the sAHP height and (G) duration. Paired t-test. (H) Representative images and quantification of SK2 and (I) SK3 expression using immunohistochemistry. Channel expression is in cyan and tdTomato expression signifying Shox2 is in magenta. SK2 and SK3 expression is lower in neonatal (P2 and P7) Shox2 INs compared to juvenile (P14) and adult (P64). Scale bars for spinal sections=200µm, insets=25µm.

### Unbiased hierarchical clustering reveals age-specific groups of Shox2 INs based on electrophysiological properties

The numerous electrophysiological properties measured in Shox2 INs shape their recruitment and ability to fire action potentials and/or bursts. In order to compare subgroupings of electrophysiologically identified Shox2 INs and possibly reveal additional age-dependent differences amongst this heterogeneous population, we next considered all 104 Shox2 neurons, spanning the three age groups, with properties measured in both current clamp and voltage clamp. We performed hierarchical cluster analysis (Garcia-Ramirez et al., 2022) using the 11 electrophysiological measurements recorded from these neurons that were not highly correlated amongst themselves. This produced 8 clusters, portrayed in the dendrogram where each leaf (individual Shox2 IN) is shaded accordingly (Fig. 6*A*). The distribution of clusters within the three age groups showed that neonatal Shox2 INs consisted only of three electrophysiological subtypes (Fig. 6*E*), clusters 1 (54%), 2 (29%), and 3 (17%), while juvenile and adult Shox2 INs consisted of seven subtypes, clusters 2-8 (Fig. 6*F, G*). Cluster numbers were ordered based on their prevalence in age-related groupings. The first three clusters, which are the only groups containing neonatal Shox2 INs, did not have a sAHP (Fig. 6*D*). Clusters 4-8, composed of only juvenile and adult cells, are characterized by similarities to cluster 3 other than the presence of sAHP (cluster 4), the presence of PICs with descending components and high input capacitance (cluster 5, Fig. 6*B*), hyperpolarized resting membrane potential (cluster 6), an increased I_M_ density (cluster 7, Fig. 6*C*), and an increased I_T_ density (cluster 8). These findings are aligned with what we have shown in Figures 2-5 by comparing these properties between the three age groups. The clustering reveals that the greatest developmental shift in the electrophysiological properties of Shox2 INs occurs following the neonatal stage, leading both juvenile and adult Shox2 INs to be relatively more heterogenous in their electrophysiology.

**Figure 6.**
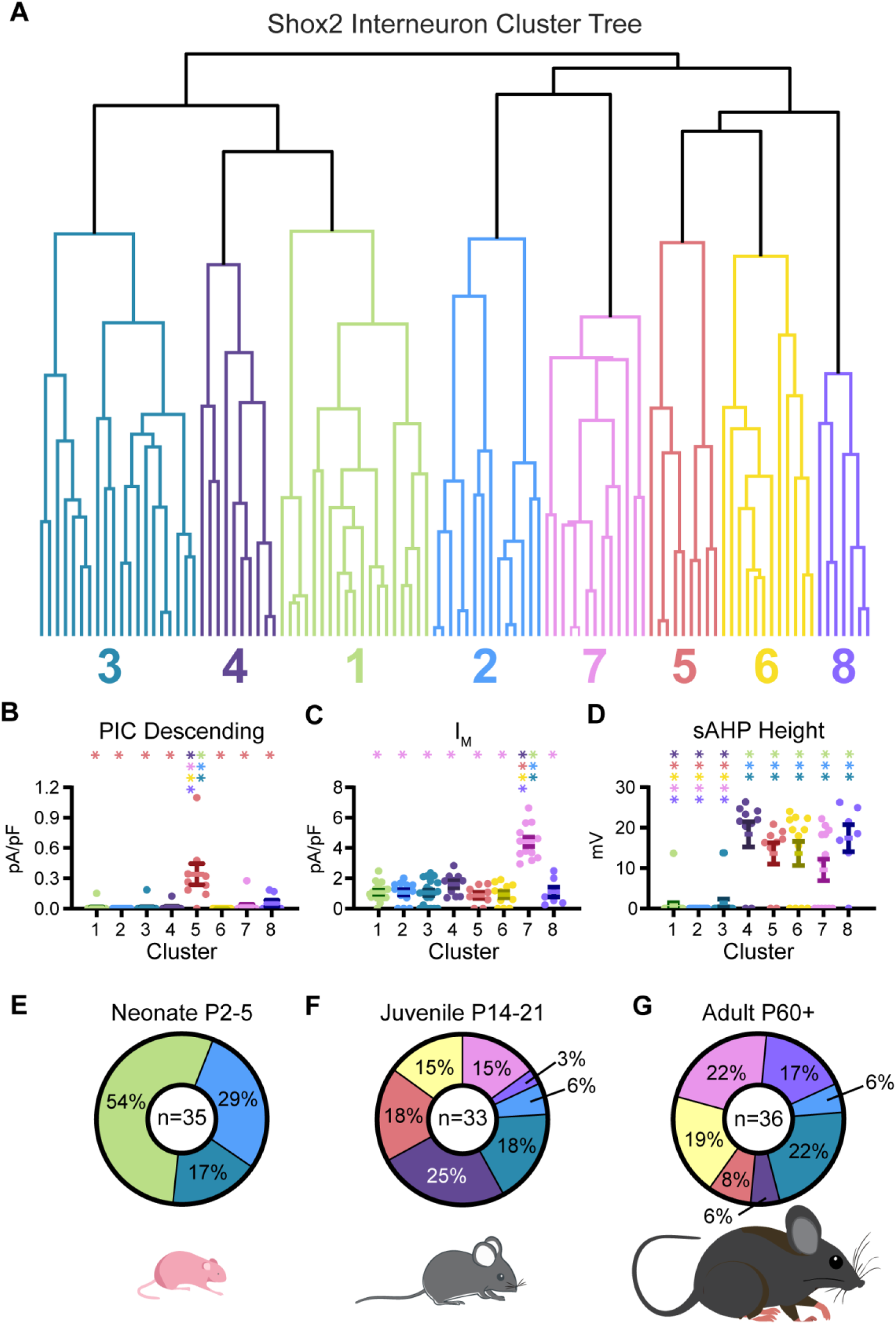
Unbiased hierarchical clustering of Shox2 Ins. (A) Dendrogram including 104 Shox2 INs, clustered according to 11 electrophysiological properties: resting membrane potential, time constant, input capacitance, threshold, sAHP height, PIC and I_H_ areas under the curve, PIC onset voltage, and current densities for PIC descending, I_T_, and I_A_. (B) Comparisons of current density of PIC descending, (C) I_M_, (D) and sAHP height between clusters, where cluster 1 has the highest percentage of neonatal Shox2 INs, and cluster 8 has the highest percentage of adult Shox2 INs. *p< 0.05, one-way ANOVA with Tukey *post-hoc* test or Kruskal-Wallis with Dunn’s *post-hoc* test. Error bars indicate SD. * Significantly different from the corresponding color-coded group. (E) Neonatal Shox2 INs are comprised of cluster 1 and subsets of clusters 2 and 3. (F) Juvenile and (G) adult Shox2 INs are comprised of subsets of clusters 2-8.

## Discussion

We investigated the changes in electrophysiological profiles and underlying channel expression within rhythm generating neurons in the lumbar spinal cord during postnatal development. Our results show that subsets of lumbar spinal Shox2 INs display numerous voltage-dependent conductances and their respective channels or RNA transcripts from neonates, juveniles capable of weight-supported stepping, and adult mice. We show that currents linked to neuronal bursting, including PIC, I_M_, and sAHP, increase in prevalence and/or magnitude as the mouse develops. Further, the diversity of electrophysiologically-defined subtypes of Shox2 INs increases with age, and subtypes of mature Shox2 INs are distinguished by the expression of voltage-gated conductances that support neuronal rhythmicity. Thus, the data suggests that the mechanisms by which Shox2 INs sustain the locomotor rhythm throughout postnatal maturation are dynamic.

### Rhythmogenic currents gain strength with age

We report that the prevalences of RNA in Shox2 INs for Na_V_1.6 and Ca_V_1.3 and the prevalences of K_V_7.2/3 in Shox2 INs do not change developmentally. The lack of change in RNA expression was surprising because it has been shown that voltage-gated ion channel expression does increase developmentally by P18 in motor neurons (Jiang, et al., 1999b). However, our data is consistent with the lack of change in the prevalences of currents with age. We were limited in our ability to characterize the membrane expression of channels which could contribute to PICs in Shox2 INs. This is because the concentrations of channels within axon initial segments in the case of Na_V_1.6 (Drouillas et al., 2023) or distal dendrites for Ca_V_1.3 (Ballou et al., 2006; Carlin et al., 2000) are difficult to detect in Shox2 neurons as they do not have a common orientation and levels of tdTomato marker labeling were low beyond the soma. Our aim in this study was to show that Shox2 INs express channels that are responsible for both sodium and calcium components of PICs observed with the electrophysiology results. We restricted our channel-specific examinations to the channel subunits most likely to contribute to PICs (Drouillas et al., 2023; Jiang, et al., 1999b; Katz et al., 2018). However, it is likely that there are other specific voltage-gated ion channels contributing to PICs in these neurons. These include Na_V_1.1 (Drouillas et al., 2023) and Ca_V_1.2 (Jiang, et al., 1999b) for I_NaP_ and I_Ca,L_, respectively. Further studies with targeted pharmacology and histology will reveal the extent to which combinations of these channels comprise PICs in Shox2 INs. In this study, we saw modest attenuation of PICs in Shox2 INs after application of riluzole or nimodipine, which is not directly compatible with our RNAscope finding of high prevalences of Na_V_1.6 RNA^+^ and Ca_V_1.3 RNA^+^ Shox2 INs. However, it is plausible that not all RNA transcripts undergo translation into proteins and that not all proteins are then trafficked to the neuronal membrane where they could contribute to the PICs measured and manipulated in this study (Solé & Tamkun, 2020). Although RNA levels are consistent, the developmental increases in PIC and I_M_ magnitudes may be caused by changes in the localization and/or density of the respective voltage-gated ion channels on the Shox2 IN membrane with age. Future studies aimed at the precise identities and locations of these channels may uncover how these intrinsic properties develop with finer precision.

In the cases of PICs which persist onto the descending phase of the voltage ramp and sAHP, the developmental shifts are distinct in that these properties are largely absent from Shox2 INs from neonatal mice. The increase in PIC magnitude with development matches the higher prevalence of PICs which persist onto the descending phase of the voltage ramp in juvenile and adult age groups, as a larger amplitude PIC is more likely to persist longer during the voltage ramp protocol. This shift may too be explained by higher PIC-related channel expression in the membrane with age, which has yet to be examined and cannot be assessed by labeling RNA. It may also be explained by a change in channel composition contributing to PICs, as other channels such as Na_V_1.1 and Ca_V_1.2 may differentially contribute to PICs in Shox2 INs throughout development. In the case of sAHP, it is clear that the neonatal Shox2 INs do not yet express SK2 and SK3. The shape of action potentials including the afterhyperpolarization is known to dramatically change with age due to changes in voltage-gated ion channels (Abbinanti et al., 2012). This has been shown across development in V2a INs (Husch et al., 2015), which partially overlap with the Shox2 population (Dougherty et al., 2013). Furthermore, neonatal V2a INs have been shown to lack biphasic AHP compared to adult V2a INs (Husch et al., 2015). In Shox2 INs this biphasic AHP appears at P10, a timepoint which may be consistent with the maturation of descending input to CPG neurons (Ballion et al., 2002; Pearlstein, 2013; Vinay et al., 2002) and is well-aligned with the establishment of weight-bearing locomotion and maturation of gait (Jiang, et al., 1999a).

### Potential to sculpt rhythmic firing behavior

To date, examinations of identified spinal IN types that extend into adulthood are limited. We show that voltage-sensitive currents in Shox2 INs gain strength with age, and that Shox2 INs from juvenile and adult mice are more heterogenous than those from neonatal mice based on their electrophysiological properties. This suggests that the intrinsic mechanisms employed by Shox2 INs to shape their function are dynamic across development. Both I_T_ and I_A_ may play more pronounced roles in Shox2 IN development. However, these currents are difficult to isolate without pharmacology due to their similar voltage-dependent properties (Huguenard et al., 1991). It is known that specific ionic conductances contribute to rhythmogenic neuronal oscillations in many CPGs. Sodium-mediated PICs (Zhong et al., 2007; Ziskind-Conhaim et al., 2008), I_T_ (Anderson et al., 2012; Ziskind-Conhaim et al., 2008), and I_H_ (Chalif et al., 2022) have been deemed essential for pharmacologically-evoked fictive locomotor activity to occur in the neonatal spinal cord, and I_M_ has been shown to alter locomotion in juvenile rats such that their speed is significantly limited (Verneuil et al., 2020). While I_T_ is only found in a subset of Shox2 INs from adult mice, it may still play a pronounced role in the rhythmogenic subset of Shox2 INs in adulthood. Interestingly, in reduced spinal cord preparations from P11-P12 mice, an age where evoking coordinated motor activity in vitro is difficult, blockade of SK-mediated conductances promotes fictive locomotor activity (Mahrous & Elbasiouny, 2018). This suggests that the appearance of sAHP in a subset of mature Shox2 INs is not directly oscillation-promoting, but that it may play a more nuanced role in shaping Shox2 IN firing behavior in that subset.

In Shox2 INs from adult mice, it is still unclear which combination(s) of electrophysiological properties are essential for rhythmogenesis. It is likely that there are many overlapping combinations of the conductances included in this study that may each lead to rhythmic firing in Shox2 INs within different contexts of locomotor behaviors. This has been thoroughly described in identified neuron types within other organisms and contexts (Alonso & Marder, 2020). The clustering we report in this study begins to highlight distinct features of mature Shox2 INs from juvenile and adult postnatal ages in which locomotor circuitry and behavior is mature. Clusters 4-8, entirely comprised of juvenile and adult Shox2 INs, are characterized by the presence of slowly inactivating PICs observed in the descending phase of the ramp, the presence of sAHP, and relatively increased current densities. This suggests that these properties may play a role in the mature functioning of Shox2 INs within locomotor circuitry. Furthermore, the post-neonatal increase in heterogeneity signifies that the processes promoting rhythmic bursting in subsets of these mature Shox2 INs are likely dynamic. The subset of Shox2 INs linked to rhythm generation (Chx10^-^) in the neonatal mouse is about 25% of the Shox2 IN population (Dougherty et al., 2013), and whether similarly sized clusters or current prevalences correspond to this subset remains to be determined. Whether the population size or electrophysiological profile of this rhythmogenic subset expands in the adult animal requires further experimentation to elucidate. Nevertheless, it is likely that mature Shox2 INs utilize the voltage-sensitive conductances, in some combination(s), including those found in this study, during rhythmic locomotor activity. Further studies are necessary to pinpoint how each of these voltage-sensitive properties can shape, mediate, or dampen rhythmic firing in mature Shox2 INs.

## Acknowledgements

We thank members of the Marion Murray Spinal Cord Research Center and Drexel ULAR for support, and we thank Ron Harris-Warrick, Jenna McGrath, Arron Hall, and Leonardo Garcia-Ramirez for comments on an earlier version of the manuscript.

